# Formation of perinuclear actin structures is driven by mechanosensitivity of actin polymerization

**DOI:** 10.64898/2026.11.09.762803

**Authors:** Berin Becic, Stephan Gekle

## Abstract

Actin filaments are one of the most dynamic components of the cytoskeleton. Their fast polymerization, particularly in regions around the nucleus, allows the cell to react quickly and efficiently to external mechanical stimuli by controling nucleus morphology and position, thereby influencing the chromatin organization and gene transcription. While the biochemical mechanisms of actin polymerization are well known, the question why polymerization happens most prominently in the perinuclear region has so far remained elusive. Here, we develop a computational model for actin network formation whose key ingredients are: (i) a fully resolved orientational distribution of the local filaments, (ii) a strain-dependent depolymerization rate and (iii) active stress generation by actomyosin motors. With only these ingredients and without any ad-hoc assumptions on the spatial distribution of signaling molecules, our physics-based model explains the formation of perinuclear actin structures after mechanical stimulation as observed experimentally. In addition, our model also provides some insight into the role actin plays in positioning the nucleus, as it predicts the occurrence of a re-centering force when the cell nucleus is displaced, in quantitative agreement with experimental observations.

## I. INTRODUCTION

The mechanics of the nucleus plays a central role for cell survivial and function. Mechanical forces exerted on the nucleus can have a decisive influence on cell fate and behavior, e.g. by influencing gene transcription [1–3], blocking multipotency [4] or inducing nuclear envelope rupture which leads to genome instability [5]. As a consequence, protecting the nucleus from unwanted deformation by external forces is a prerequisite for long-term stability of cell function [6–11]. On the other hand, actively generated intracellular forces enable the cell to control the nucleus position and shape which appears to be a key determinant for cell motility in particular in 3D environments [12].

The two key structures for force transmission to the nucleus that have emerged from experimental work of the last decades are the so-called perinuclear actin ring [11, 13] and the nuclear actin cap [7, 11, 14, 15]. In addition to experimental studies, computational models have been developed [7, 16, 17] to capture the mechanical coupling between the cell exterior, the cytoskeleton and the nucleus. These latter studies focussed mostly on the effects of existing actin structures around the nucleus, but did not attempt to clarify in detail the mechanisms which lead to their formation in the first place. For example, it has been observed that the perinuclear actin ring assembles very quickly (∼ 20*s*) upon sudden mechanical stimulation but it is unclear what drives this fast and highly localized assembly [13, 18].

Here, we present a single computational model which encompasses both the formation and the functioning of the perinuclear actin ring. We apply our model to two different prototypical experimental setups. First, our model describes a biophysical mechanism for the formation of an actin ring around the nucleus after a mechanical stimulation of the cell [13, 18]. Second, we explain the re-centering of a displaced nucleus by the formation of an inhomogeneous actin ring which exerts a force on the nucleus directed towards the center [19].

Our starting point are well-established continuum active gel models which have been very successful in capturing a variety of different aspects of the actin cytoskeleton [20–22]. These typically describe the actin network either as isotropic using a scalar concentration field or account for its directionality by using a nematic orientation parameter which describes the local mean orientation of the actin filaments. A series of different works, starting from the one by Deshpande et al. [23], used a more general approach and included a spatially dependent orientation distribution of actin filaments [24–26]. With this, they were able to observe ring-like structures around the nucleus even though they did not study them further [24, 27]. Here, we develop a generic, microscopic model for the polymerization and active stress generation of actin filaments around the nucleus which allows us to explain the actin dynamics for the two prototypical experimental situations introduced above.

## II. METHODS

### A. Biophysical model

#### 1. Continuum hypothesis

The actin structures which we aim to describe consist of individual actin filaments which are cross-linked by a variety of actin binding proteins. For branched actin networks like the actin cortex or the lamellipodium the length of an individual actin filament lies between 100 − 1000*nm* [28–30] whereas in stress-fibers they tend to be longer and reach up to ≈ 2*µm* [31]. As we are interested to model the actin network on the length scale of the cell (∼ 30*µm*) we use a continuum approach and introduce the following two quantities: The number density of actin filaments per angle is denoted as 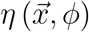 and the number of monomers in one actin filament with the respective orientation is denoted as 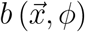. A crucial feature of this description is that these fields not only depend on the spatial coordinate 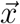 but also on the angle *ϕ* describing the orientation of the filaments. Such an approach was first introduced by Deshpande et al.[23] and later modified [24–26]. Here, we implicitly assume that the cell is flat (see next section) which is the case for adherent cells. As a result the actin filaments are all oriented approximately in the horizontal plane (parallel to the substrate) and their orientation in that plane can be described by one single angle *ϕ*. Thus, the number of actin filaments in the volume 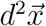 around 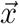 and an orientation between *ϕ* and *ϕ* + *dϕ* is 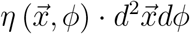. The total number density of actin monomers which have polymerized into the network in this volume is 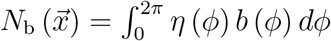.

The dependency of these fields on the angle *ϕ* describes the geometry of the resulting actin network. For fully aligned filaments, 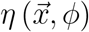 is high only along the fiber direction, i.e. 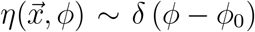 with *ϕ*_0_ the fiber direction. In contrast, for an isotropic network the angular density is constant, i.e. 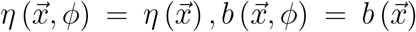. Additionally, our description implicitly assumes that all actin filaments at one position which have the same orientation also have the same length. We imagine that this is enforced by the geometric constraints posed by the cross-linking proteins.

To continue, we further assume that no transport of actin filaments occurs. We therefore only describe the incorporation of cytosolic actin filaments into the network. These actin filaments form by the continuous polymerization of actin monomers in the cytosol. This assumption is motivated by the experimental finding that the formation of the perinuclear actin ring is driven by actin polymerization in the vicinity of the nucleus [13], as well as the immobilization of different cell organelles during the formation of the actin ring [18].

**FIG. 1:**
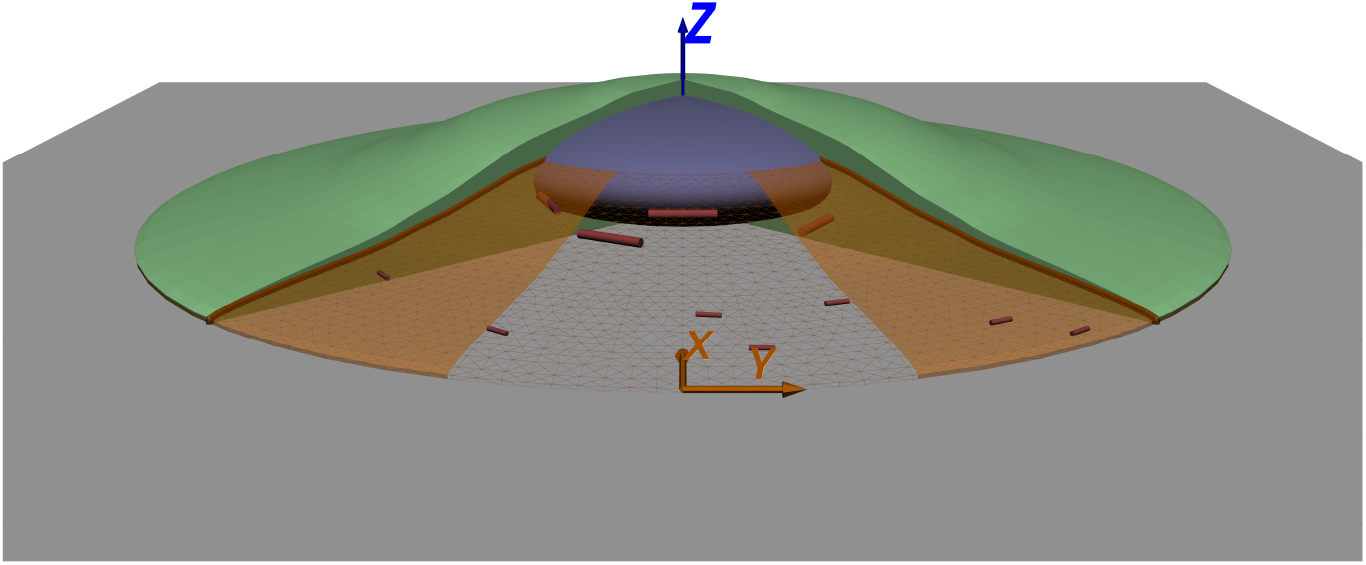
Illustration of an adhered, relatively flat 3D cell with the cytoskeleton in green and the nucleus in blue. The slice described by our 2D model (orange) starts at the outer edge and follows the cell contour just below the upper surface until it reaches the nucleus.

#### 2. Cell geometry and passive elasticity

In agreement with the experimental situations that we attempt to model, we consider a strongly adhering cell whose height is small compared to its lateral extension. Accordingly, we use a two-dimensional description to reduce the computational effort [23, 32, 33]. In three dimensions, the height of the cell is not constant, but decreases towards the edges, described by an elevation profile *h* (*x, y*). To account for this we consider a two-dimensional slice following the elevation profile of the cell 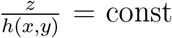 rather than a plane at a fixed *z* = const as is typically done in 2D formulations. Such a slice is sketched in brown in figure 1. To achieve the best representation, we choose a slice which is relatively close to the upper cell surface but still sufficiently intersects the nucleus. At the edge of the cell, the slice coincides with the substrate to which the cell firmly adheres and one therefore obtains the boundary condition 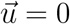 with the displacement 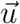.

Cytoskeletal components other than actin filaments are considered as a passive elastic continuum and their stresses are modeled using linear elasticity:

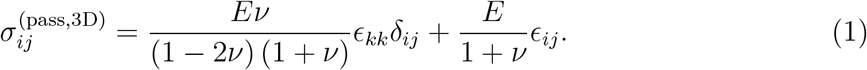

Here, *E* denotes the elastic modulus and *v* the Poisson ratio. As usual, *ϵ*_*ij*_ represents the displacement gradient tensor. We apply the free force boundary condition *σ*_*ij*_*n*_*j*_ = 0 at the upper cell surface.

As the cell is very thin, 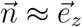 this leads to 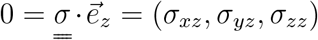 and thereby implies 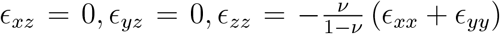. By inserting this back into the three-dimensional stress tensor we obtain the following passive stress tensor:

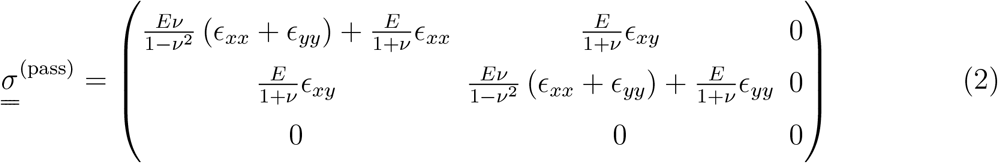

with all *z*-components equal zero, such that in the following we only consider the *xx, xy, yx* and *yy* component. As reference stress-free state for the passive elasticity, we assume the initial shape.

#### 3. Time evolution of the actin network

To model the time evolution of the actin network we consider two processes: The growth of the network via the incorporation of new actin filaments and the remodeling, i.e. the incorporation of new actin monomers into already bound filaments. Both processes heavily depend on a wide variety of proteins and signaling molecules. As we focus on the mechanosensitive aspects and how they control the nucleus we will for simplicity assume that the biochemical environment is temporarily and spatially constant and favors the polymerization of the actin network. Consequently, the state when an actin filament is bound in the actin network is energetically favorable compared to the unbound state.

##### a. Incorporation of new actin filaments

To start, we consider the growth of the actin network via the incorporation of new actin filaments, as illustrated in figure 2a. Here we assume that once an actin filament with the same number of monomers as those already bound in the network and suitable cross-linkers simultaneously meet at the network they immediately bind to it. This leads to a constant binding rate per unbound filament *ω*_1_ which is limited by the diffusion and the availability of these constituents in the surrounding cytosol. We do not explicitly consider the processes leading to the growth of the actin filaments from the monomeric actin pool in the cytosol, as it is much faster than the incorporation of the filaments into the network. Instead, we estimate the number of suitable actin filaments which are available for incorporation into the network at any given time to be 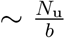 where *N*_u_ denotes the number density of available actin monomers which we assume to be constant throughout the cell as detailed below. This leads to the polymerization rate of the actin network 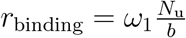.

**FIG. 2:**
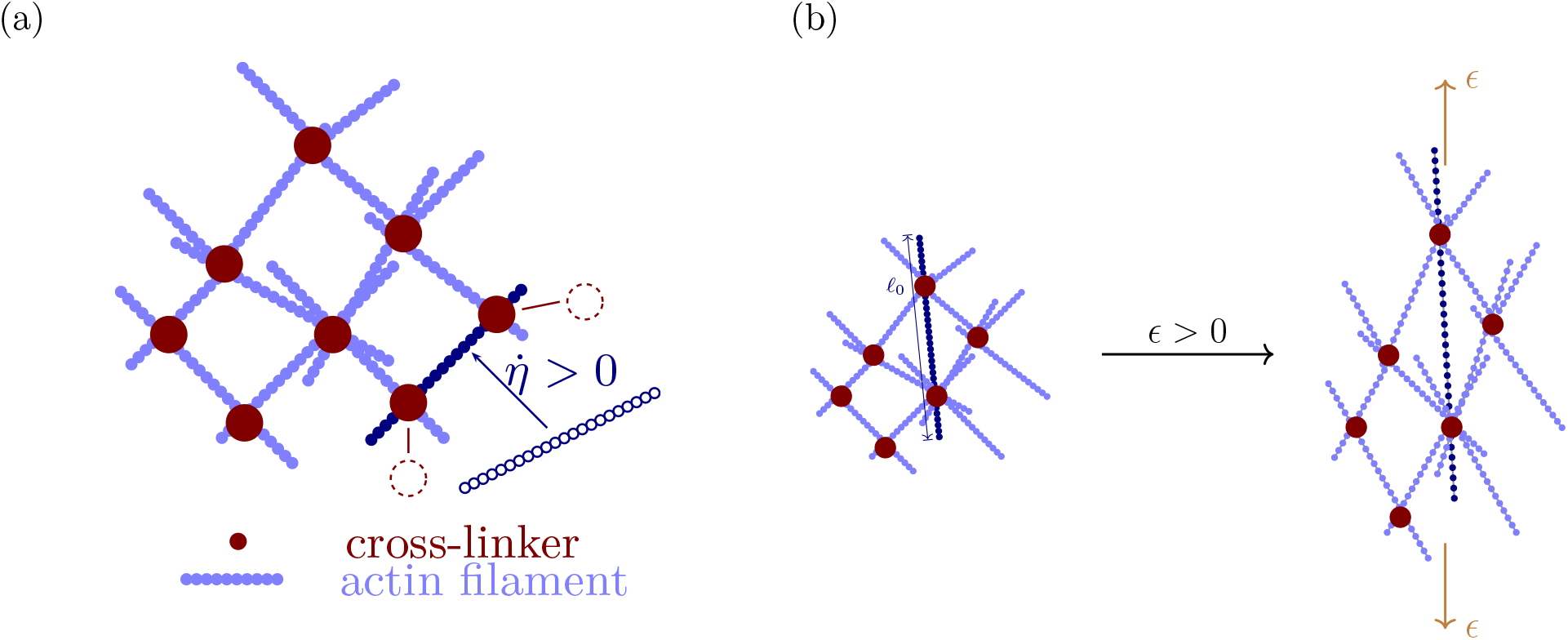
Illustration of the two basic mechanisms for actin dynamics in our model. (a) sketches the incorporation of new actin filaments in the network (polymerization). (b) illustrates the change of the filament length upon the stretching of the network (remodeling).

Next we consider the unbinding process of an actin filament from the network. As this involves the breaking of the cross-linking bonds, the actin filaments have to overcome an energetic barrier Δ*µ* > 0 leading to a depolymerization rate per filament 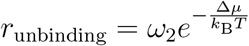. As the number of cross-links on an individual actin filament increases with the number of monomers it is reasonable to assume that this energy barrier increases linearly with its number of monomers 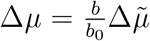. Here, *b*_0_ denotes the number of monomers in a fully relaxed actin filament in an undeformed cell. The undeformed state is specified by the reference state of the passive elasticity described in section II A 2. Consequently, 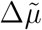 is the binding energy of an actin filament containing *b*_0_ monomers. Then, the time evolution of the concentration of bound actin filaments can be summarized to:

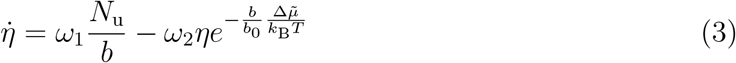

Finally, we have to determine the density of unbound actin monomers *N*_u_. We assume that their concentration is spatially constant, as in contrast to the bound actin network the small monomers can diffuse quickly through the cell cytosol. As typical diffusion constants are 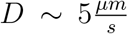 [34, 35] and the actin ring builds up over *t* ∼ 40*s*, monomers diffuse by 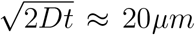 which corresponds to the typical distance between the periphery and the nucleus in the situations considered here [13, 18]. For the re-centering considered in section III B the time scale of the evolution of the actin structures is even larger. Nevertheless, we did check that allowing a spatially dependent *N*_u_, i.e. neglecting actin monomer diffusion, does not significantly affect our results. The total number density of (bound and unbound) actin monomers can thus be written as

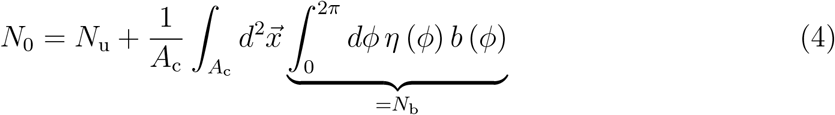

where *N*_0_ depends neither on time nor space, *N*_u_ depends on time only and *N*_b_ depends on time and space. The cell area excluding the nucleus, where no actin is present, is denoted *A*_c_. Since we consider small deformation only, *A*_c_ and coordinates 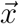 always refer to the undeformed shape.

##### b. Remodeling of bound actin filaments

Next we describe the remodeling of the actin filaments in the network. To start, we consider an undeformed cell which corresponds to the reference state of the passive elasticity discussed in section II A 2 and an isotropic actin network which is formed in it. We define *l*_0_ as the physical length and *b*_0_ as the number of actin monomers, respectively, in one filament in this state. These quantities will, in general, strongly depend on actin regulating proteins which here are considered to be spatially and temporally constant.

Now, figure 2b illustrates the stretchig of the entire cell from *l*_0_ to *l*^*′*^ = *l*_0_ (1 + *ϵ*) which increases the distance between monomers away from the relaxed state distance *b*_0_/*l*_0_. The filament will react by incorporating new monomers until the new equilibrium *b*_eq_ is reached which fulfills 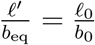. Inserting *l*^*′*^ from above, the new equilibrium is given by *b*_eq_ = *b*_0_ (1 + *ϵ*) with *ϵ* = *ϵ*_*ij*_*n*_*i*_*n*_*j*_ being the component of the strain tensor *ϵ*_*ij*_ in filament direction. Denoting the filament orientation vector as 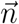, this strain component is *ϵ* (*ϕ*) = *ϵ*_*ij*_*n*_*i*_*n*_*j*_. To describe the dynamics of this incorporation, a very simple model is employed, assuming that the filaments relax exponentially towards their new equilibrium:

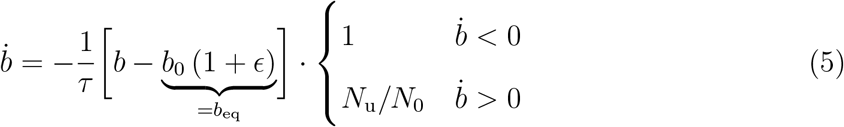

Here *τ* is the timescale of the remodeling process and the prefactor *N*_u_/*N*_0_ accounts for the effect that the incorporation of new monomers is limited by the availability of actin monomers in the surrounding cytosol.

To sum up, equations (3), (4) and (5) govern the time evolution of the actin filament concentration and their number of monomers for every orientation independently.

##### c. Steady state

To interpret the mechano-sensitivity of the actin network formation resulting from this model, we determine the steady-state concentration of bound actin monomers *N*_b_ for a uniaxial cell strain *ϵ*_*yy*_ which, without loss of generality, is chosen along the *y*-direction. Note that this leads to an orientation-dependent scalar strain *ϵ*(*ϕ*) which is maximal in the *y*-direction. The concentration of bound actin monomers is obtained from the previously derived time evolution equations of the actin network. Setting 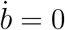 in equation (5), we obtain *b* = *b*_eq_ = *b*_0_ (1 + *ϵ*) while setting 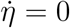 in equation (3) yields

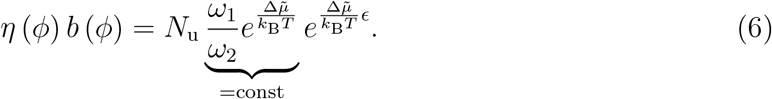

Thus, for small concentrations of bound actin monomers where *N*_u_ ≈ *N*_0_ = const, the amount of bound actin monomers *η*·*b* grows exponentially with *ϵ*_*yy*_. For large concentrations, the growth of *η* · *b* slows down as *N*_u_ → 0. This behavior is illustrated in figure 3a where the dependency of the total concentration of bound actin monomers on the uniaxial strain *ϵ*_*yy*_ is plotted. For *ϵ*_*yy*_ = 0, we obtain the reference concentration in the unstressed state 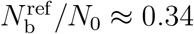.

**FIG. 3:**
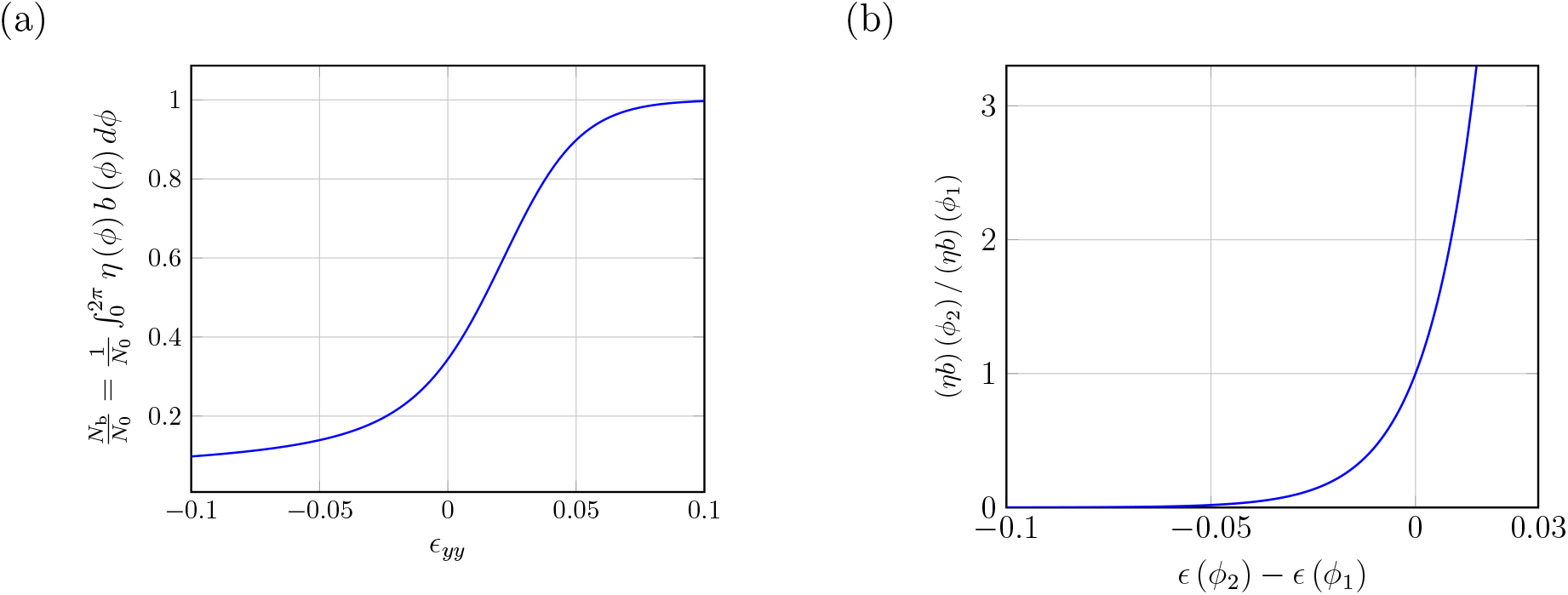
Concentration and stress generation of the actin network in the steady state. (a) shows the dependency of the total concentration of bound monomers *N*_b_ on a uniaxial cell strain *ϵ*_*yy*_. (b) shows the exponential dependency of the ratio of bound actin monomers in different orientations on the difference of the respective strains.

Next, we consider anisotropic cell strain. Then, the steady-state concentration of filaments in each direction will depend on their orientation relative to the cell strain. Interestingly, our model predicts that the ratio of bound actin filaments with different orientations *ϕ*_1_, *ϕ*_2_ increases exponentially with their strain difference

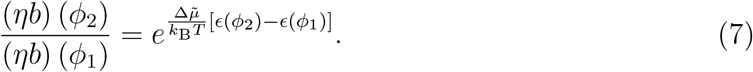

This implies that a static anisotropic strain leads to an alignment of the filaments in the direction of the maximal extensile strain as illustrated in figure 3b. These strain-dependencies can be qualitatively understood as follows: If the cell contracts (*ϵ* < 0) the actin filaments become compressed. As they remodel towards their relaxed state following equation (5) they lose actin monomers and thus also the number of cross links decreases. As a result, the total binding energy of the actin filament to the network decreases which finally leads to a higher dissociation rate and a lower concentration of bound actin in the steady state. We mention already here that a contraction of the cell can be caused by acto-myosin contraction which again depends on the actin network. This feedback will become crucial later to understand the formation of the actin structures in section III A.

#### 4. Active stress generation

To complete the model we now specify the active stress generated by the acto-myosin network. Active stresses arise when myosin complexes bind to two neighboring actin filaments, undergo a conformational change and then pull the actin filaments together [28]. As a result, the tension *T* depends on the actin network and thus the fields 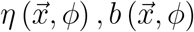 and can be derived as follows. In our continuum model, all quantities including *T* are averages over a volume element which here is assumed to be small enough that maximally one myosin inside the volume is in its force generating state at a certain time. Then, the average tension scales linearly with the time fraction in which one myosin complex is in its force generating state. This fraction, in turn, depends linearly on the total number of myosin binding sites which scales linearly with the number of actin monomers bound in the network. In total, the tension generated by filaments with orientation 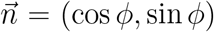 scales as *T* ∼ *η*(*ϕ*) · *b*(*ϕ*).

Denoting as *T*_max_ the tension which is achieved when all actin polymerizes the tension per actin monomer is 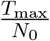 and thus

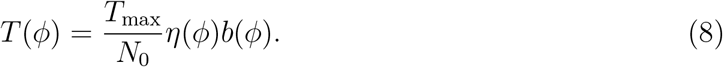

From this, the full three-dimensional stress tensor *σ*_*ij*_ is obtained by integrating *T* over all directions *ϕ* and adding the passive stress 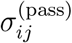 from equation (2):

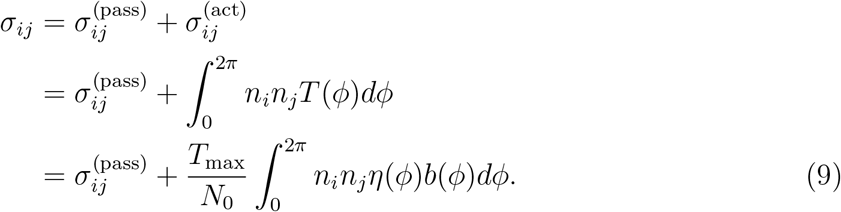

This stress tensor combined with the condition for a quasistatic force equilibrium

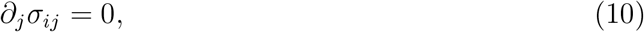

together with the time evolution equations for the actin network (equations (3), (4) and (5)) fully specify our model in the cytoskeleton region outside the nucleus.

#### 5. Model for the nucleus

Experimental data suggests that the nucleus behaves as a purely elastic material under physiological strains [36, 37]. Its elastic modulus is, however, not reliably known and is most likely not constant, since nuclear deformation might affect the cross linking of lamin proteins and chromatin organization increasing nuclear stiffness [38, 39]. As we are primarily interested in the influence of the actoymosin network on the nucleus we choose its elastic properties to be the same as the passive elastic properties of the rest of the cell. This implies that the deformation and normal stresses are continuous at the outer boundary of the nucleus. We found that the effects which we observe in this work do not change qualitatively when varying the nucleus’ elastic modulus.

### B. Parameter choice

In this section, the choice for the parameters employed in the following simulations will be presented and justified. To start, we note that the governing equations ((3), (4) and (5)) only depend on the ratios 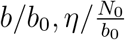 and *N*_u_/*N*_0_. Thus, the quantities *N*_0_, *b*_0_ can be eliminated by normalizing *N*_u_, *b, η* accordingly and consequently do not need to be specified (see appendix A for the resulting equations). Furthermore, they do not explicitly depend on the cell size but only on the ratio *R*_nucleus_/*R*_cell_ which is set to 0.4. The values of all remaining parameters are listed in table I and will now be justified.

**TABLE I:** This table lists all the parameters of the model including the chosen values. A detailed justification for these choices is given in the text.

|  | **active and passive stresses** |  |
|---|---|---|
| *T*<sub>max</sub> | maximum acto-myosin tension | 4*kPa* |
| *E* | passive elastic modulus | 500*Pa* |
| *ν* | Poisson ratio | 0.45 |
|  | **actin network dynamics** |  |
| Δ*μ̃* | dimensionless chemical potential difference of the unbound and bound state per monomer | 80*k*<sub>*B*</sub>*T* |
| *ω*<sub>1</sub> | normalized binding rate | 0.05*s*<sup>−1</sup> |
| $\omega_2 e^{-\frac{\Delta\tilde{\mu}}{k_B T}}$ | unbinding rate | 0.61*s*<sup>−1</sup> |
| *τ* | relaxation time of the actin filament remodeling process | 3.3*s* |

As passive elastic parameters we choose *E* = 500Pa, *v* = 0.45. The measured elastic response of cells varies heavily depending on the cell type, substrate and measurement method, but typical values lie around *E* = 1kPa [40]. These measurements, however, represent overall values that do not separate the passive elastic response from the contribution of the acto-myosin network as we do in our model. To estimate only the former, we consider experiments [41, 42] which showed that the inhibition of the acto-myosin contractility significantly reduces the elastic modulus to around 30 − 50% of its initial value. We therefore use a smaller elastic modulus of *E* = 500Pa for the passive contribution in our model. Next, we set the maximum contractile tension *T*_max_ = 4kPa. This is based on measurements of the contractile tensions occurring in focal adhesions [43] which we expect to be close to the maximally achievable acto-myosin tensions in non-muscle cells.

For the dynamics of the actin network, in general, no direct experimental values are known such that only rough estimations can be made here. The difference of the chemical potential of the free and bound state 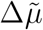 determines the strain sensitivity of the actin network and thus the extent of the actin ring. We estimate 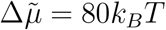 based on the binding energy of the cross links between two actin filaments. From molecular dynamics simulations binding energies per cross-linking molecule of around 10k_*B*_*T* are reported [44–46]. The mesh size in cortical actin which lies around 20*nm* delivers a rough approximation for the distance between individual cross-linking sites. Individual actin filaments are ∼ 200*nm* long [47] and thus contain ∼ 10 cross links which implies a total binding energy per actin filament in the order of magnitude 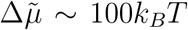. As the maximal strains occurring are ∼ 0.1, the binding energy between an unstrained and a maximally strained filament differs maximally by 10k_*B*_*T*.

Next, the binding (polymerization) and unbinding (depolymerization) rates per filament are set as *ω*_1_ = 0.05*s*^−1^ and 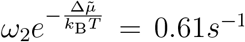. Assuming that the growth of the actin network by the incorporation of new actin monomers is the slowest time scale, the depolymerization rate 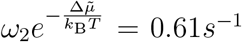 was fitted such that the model reproduced the experimental time scale for the occurrence of the perinuclear actin ring which is around 40*s* [13]. Subsequently, the polymerization rate *ω*_1_ was fitted such that the extent of the actin ring becomes maximal.

Finally, the timescale for the remodeling of the actin network is set to *τ* = 3.3*s*. This choice is motivated by the turnover timescales of the actin network which are known to be strongly influenced by the presence of several actin-binding proteins and vary from seconds in cortical actin and the lamellipodium up to hours for permanent structures like sarcomeres [48, 49]. As the actin structures considered here are rather disorganized we expect that their turnover times are similar to those in dynamic structures like the actin cortex and the lamellopodium, justifying the choice *τ* = 3.3*s*. As a result, the remodeling is the significantly faster process. I.e. the incorporation of new filaments (*ω*_1_, *ω*_2_) is the limiting factor determining the timescale of the build-up of the actin ring, whereas the time scale of the remodeling (*τ*) only weakly influences how fast the actin ring develops.

### C. Numerical method

As we consider a two-dimensional domain, we discretized the cell (including the nucleus) using triangular linear elements. Typical resolutions for most of the simulations were *N*_triangles_ = 10000. The governing equations were solved using a homemade finite-element code. Directly around the nucleus a higher resolution was used (∼ 4*×* as high) to accurately resolve the actin ring. The displacement 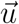 was interpolated linearly over these elements while the fields *η* and *b* are considered constant within each triangle. Besides the spatial discretization, we discretized the orientational dependency of *η* and *b* using *N*_*ϕ*_ = 200 points.

The orientational integrals were evaluated using the trapezoidal rule

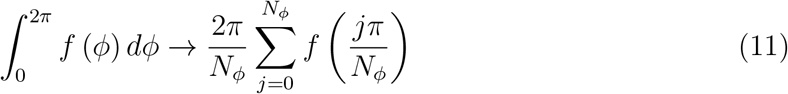

where additionally the symmetry *ϕ* ↔ *ϕ* + *π* was exploited.

For time stepping, the following scheme was used: Based on the current displacement and the resulting strain, for each triangle, the evolution equations for the actin network, (3) and (5), define a set of 2*N*_*ϕ*_ ordinary differential equations which are independent for each triangle. Thus for each triangle the corresponding set of evolution equations was integrated using the Cash-Karp method. The stress tensor depends on the strain which can be expressed using the new displacements 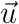 (*t* + Δ*t*) and the strain-rate which is also expressed in terms of the new displacements 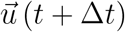 using forward differences: 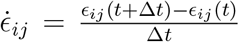. Using the condition for mechanic equilibrium *∂*_*j*_*σ*_*ij*_ = 0 we determined the new displacements 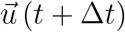. During this step the new values for the actin fields *η* and *b* were used and 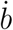 was determined using equation (5) by inserting the strain from the last time step. The nucleus was realized by not integrating the evolution equations for the triangles which make up the nucleus, implying *η* = 0 for all triangles whose initial centroid position is within *R*_nucleus_ from the nucleus center.

As initial conditions, we chose 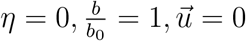.

## III. RESULTS

Using the model presented above, we aim to understand two key experimental findings. We start with the formation of the perinuclear actin ring which is considered in the subsection III A.

References [13] and [18] observed its formation upon a sudden mechanical stimulation which triggered an influx of calcium ions leading to a relatively homogeneous and instantaneous increase in the concentration of Ca^2+^-ions. These activate the actin nucleator inverted formin 2. Intriguingly, although both the nucleator and the elevated concentration of Ca^2+^-ions were present throughout the entire cell, only a highly localized actin ring around the nucleus formed. As we will show below, our model can explain this finding by the exponential dependence of actin depolymerization postulated in equation (3).

Next, we study the influence of the actin network on the positioning of the nucleus. In general, there exist a big variety of different mechanisms involved in the nucleus positioning which, next to acto-myosin contractility also include pushing forces exerted by actin polymerization or the activity of microtubule-associated motors [12]. Here, we restrict ourselves to model one particular situation: [19] studied the recentering of nuclei after their displacement by centrifugation and found that in certain situations it necessitates the binding of actin to the nucleus and acto-myosin contractility.

### A. Formation of the perinuclear actin ring

We model the experiments of Shao et al. [13] starting from the moment directly after mechanical stimulation, when calcium is present throughout the cell, but actin polymerization has not yet started. Accordingly, we use the initial condition *η* = 0 and set *ω*_1_ > 0 as explained in subsection II B. When solving our model, we reach a steady state after ≈ 40*s*. As shown in figure 4, we clearly find the formation of a perinuclear actin ring in agreement with the experimental observation. The spatial distribution of bound actin, 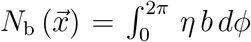, is shown in figure 4a demonstrating the ring as a thin region around the nucleus with an elevated actin concentration *N*_b_/*N*_0_ ≈ 0.42. At the outer edge of the cell a concentration of *N*_b_/*N*_0_ ≈ 0.3 is observed. Comparing with the unstressed reference concentration 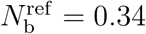 from section II A 3 c, we find that the presence of the nucleus induces actin polymerization near the nucleus and actin depolymerization in the cortex region.

**FIG. 4:**
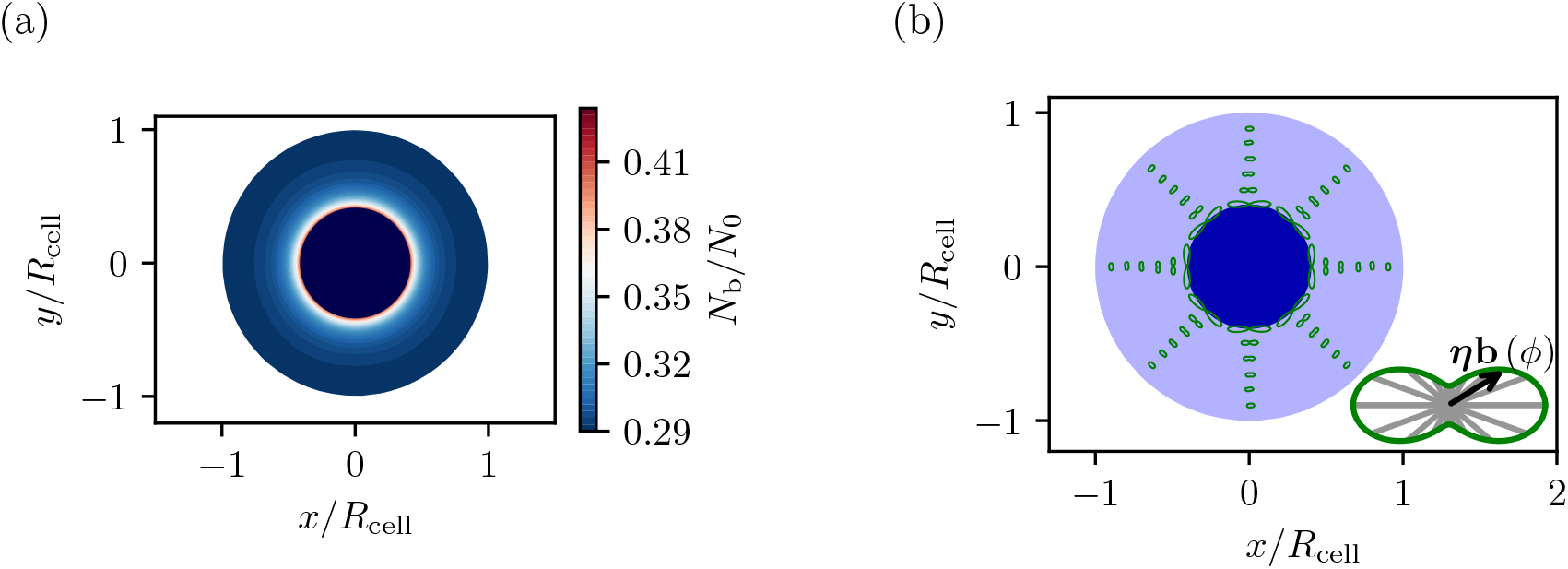
Formation of the actin ring around the nucleus after mechanical stimulation. (a) shows the concentration of bound actin monomers throughout the cell. (b) shows the orientations of the actin filaments using polar plots of *η* (*ϕ*) · *b* (*ϕ*). The length of the arrow, i.e. the distance of the green curve from the center, indicates the local concentration of filaments with the respective orientation. Both plots show the situation when the steady state is reached.

Interestingly, depolymerization of cortical actin after a mechanical stimulus has also been observed experimentally by [13, 18]. Figure 4b shows the orientation of the actin filaments. The orientation distributions (*ηb*) (*ϕ*) at different points are illustrated as polar plots. We observe that in addition to the higher concentration, the filaments in the actin ring are strongly aligned tangentially around the nucleus.

To understand both the increased actin concentration near the nucleus as well as the tangential orientation of the perinuclear filaments, we consider the steady-state anisotropic cell strain shown in figure 5a. We first note that the acto-myosin contraction in the cytoskeleton leads to an isotropic expansion of the nucleus, i.e. *ϵ*_rr_ = *ϵ*_*ϕϕ*_ > 0 for *r*/*R*_cell_ < 0.4. This is consistent with the experimentally observed expansion during the mitotic exit triggered by actin polymerization [50] as well as its shrinking after the inhibition of myosin contraction or the depolymerization of actin [51]. Next, we consider the radial strain *ϵ*_rr_ outside the nucleus for *r*/*R*_cell_ > 0.4. Since the cell adheres to the substrate at its periphery the deformation u = 0 at *r*/*R* = 1. Accordingly, the expansion of the nucleus cannot be compensated by a corresponding expansion of the cytoskeleton and therefore the latter has to undergo radial contraction which leads to *ϵ*_rr_ < 0 as shown in figure 5a. At the same time, the elongation of the nucleus perimeter causes a positive tangential strain *ϵ*_*ϕϕ*_ of the cytoskeleton around the nucleus which vanishes towards the cell boundary as a result of the adhesive boundary condition 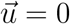. Refering back to figure 3b, this leads to an increased depolymerization rate of radially oriented filaments and a decreased depolymerization rate of tangentially oriented filaments. This finally leads to the predominantly tangential alignment of the actin filaments. Refering back to the exponential dependence of the equilibrium actin concentration on cell strain (figure 3a) we see furthermore that the concentration increase of tangential filaments outweights the concentration decrease of radial filaments thus explaining the net increase of polymerized actin in the vicinity of the nucleus as shown in figure 4a. We conclude that the exponential strain dependence of the actin depolymerization rate postulated in equation (3) can be considered as the root cause for the formation of the perinuclear actin ring.

**FIG. 5:**
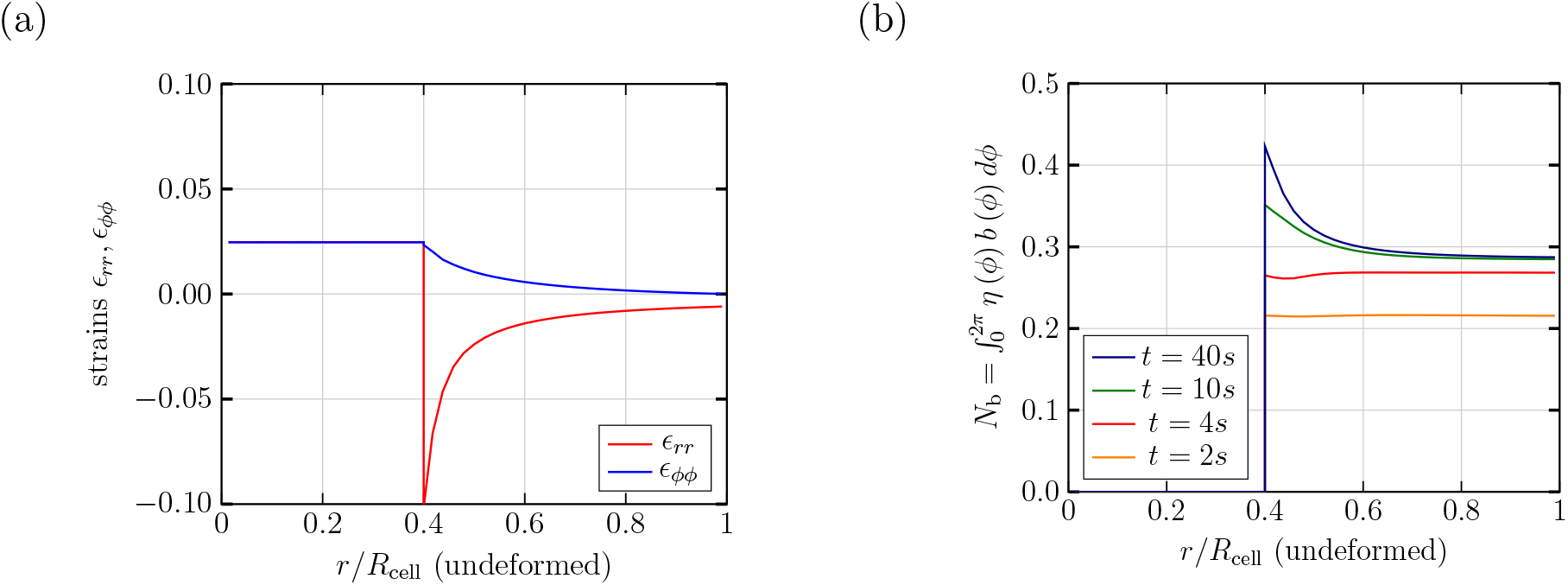
(a) shows the cell strain split into radial and tangential components for the same situation as in figure 4. (b) illustrates the time evolution of the actin ring by showing the radial profiles of the total concentration of bound actin at different times.

In our proposed mechanism, contractile actomyosin tensions play a central role for the formation of the actin ring. We therefore checked the sensitivity of the mechanism to the magnitude of the contractile stress by varying *T*_max_. In appendix B we show that even for an eight times smaller value of *T*_max_ our model still predicts formation of the actin ring. This is in agreement with the experiments in [13, 18] who found that the actin ring still occurred after adding blebbistatin which inhibits actomyosin contractile activity.

Finally, we investigate the temporal dynamics of actin ring formation in figure 5b which shows the radial actin concentration profiles at different times. We predict that there is an initial transient of about 5*s* in which the actin concentration near the nucleus is actually lower than in the periphery. However, this transient vanishes quickly and after about 40*s* the steady state as described in the previous paragraph is reached.

### B. Recentering of the nucleus

Next to controlling its shape the actin cytoskeleton is also known to play a role in positioning the nucleus, e.g. during cell migration or nucleokinesis [9]. As a second application, we use our model to gain insight into the re-centering of an initially displaced nucleus as in the experiments of Zhu et al [19]. In these experiments, the nucleus was first moved to an off-centered position by centrifuging the cells, after which it was observed how the nucleus returned to its original position ∼ 20mi*n* after the centrifugation was stopped. To mimic this situation we use a passive elastic reference shape in which the nucleus is displaced from the center by δ*y* = 0.5*R*_cell_ and leave all other boundary conditions unchanged as in section III A. The resulting distribution of bound actin and its orientation are shown in figure 6.

**FIG. 6:**
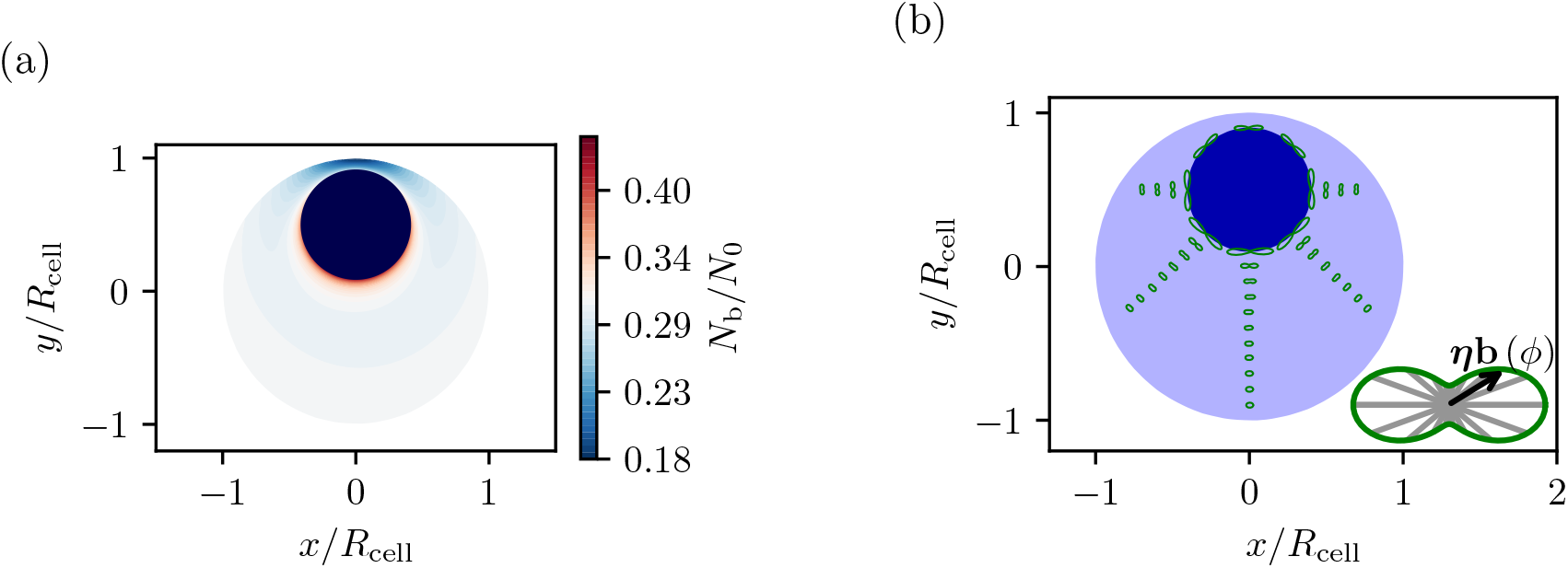
Steady-state actin network in a cell with a displaced nucleus. (a) shows the corresponding total concentration of bound actin which is higher on the side of the nucleus facing the cell center. (b) shows the orientation distribution of the actin filaments in the steady state for a cell with a displaced nucleus.

Similar to the previous cases, the actin filaments are mainly oriented tangentially. In contrast to the previous situations, however, the actin concentration on the nucleus side facing the cell center is higher than on the side facing the boundary. The reason for this difference, according to our model, is that the strain on the side closer to the cell boundary is lower due to the fixed cell edge. This lower strain leads to shorter filaments and thus to a higher depolymerization rate which reduces the concentration of actin filaments compared to the side facing the cell center. This discrepancy in concentrations leads to a discrepancy in acto-myosin induced stressess and, finally, to a force pulling the nucleus back to the cell center.

To obtain a quantitative estimate, we first calculate the force which is exerted by the acto-myosin contraction on the nucleus, by only considering the active part of the stress tensor from equation (9):

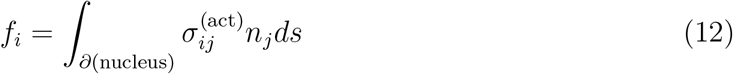

Note that as we are always only considering a two-dimensional slice of the cell f_*i*_ is a force per unit height of the cell. We use a deflection of the nucleus δ*y* = 0.05*R*_cell_, set the cell radius *R*_cell_ = 30*µm* and choose a nucleus size of *R*_nucleus_ = 0.4*R*_cell_ as in the paragraphs above. For this situation, our model predicts a total force per height on the nucleus of 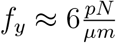 as obtained by numerically evaluating the integral in equation (12) from our simulation results. As height of the cell, we use *h* = 6*µm* which is a typical value for monolayers of the same fibroblast cell line [52] as used by Zhu et al [19] to study this recentering. To estimate the resulting velocity we assume that for very slow movements this force is compensated by the viscous drag which is very roughly ∼ 6*πR*_nucleus_*η*v_recenter_. As typical value for the static viscosity of the cytoplasm we use *µ* ≈ 60Pa · *s* as it was found for the same cell line via magnetic rotational spectroscopy [53, 54]. ^1^ With this, we finally obtain

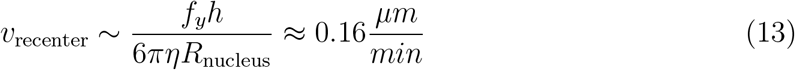

This result is close to the experimentally found recentering velocity of 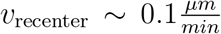 which we estimated based on figure 2d) of Ref. [19]. This final application provides further evidence for the general applicability of our model, even beyond our initial motivation to explain the formation of a static perinuclear actin ring.

## IV. CONCLUSION

We presented a theoretical model which predicts the formation of perinuclear actin structures triggered by a mechanical stimulus to biological cells, in agreement with experimental observations. The perinuclear structures exhibit an enhanced concentration of polymerized actin filaments together with a predominantly tangential filament orientation in a thin region around the nucleus. Importantly, our model does not require an enhanced concentration of nucleators or signaling molecules in the perinuclear region. Instead, the key mechanism in our model is an exponential decrease of the depolymerization rate as a function of the local mechanical strain. Together with active stress generation by myosin molecular motors bound to actin filaments, this creates a positive feedback loop leading to both enhanced concentration and tangential orientation of actin filaments in the perinuclear region. All ingredients of the model are microscopically motivated without ad-hoc assumptions and realistic, experimentally motivated values are used for all model parameters.

As a second application, we consider a cell whose nucleus is displaced from the cell center. In this situation, without any further ad-hoc assumptions, our model predicts the existence of a net force pushing the nucleus back towards the cell center. From this force, we can estimate the centripetal velocity with which the nucleus is expected to move which we find in quantitative agreement with experimental observations.

In conclusion, our model provides a generic, physics-based mechanism for the formation of localized actin structures without the need of specific spatial distributions of biochemical signaling molecules.

## Appendix A: Non-dimensional governing equations

Here the non-dimensionalized equations for the evolution of the actin fields and the stress tensor are given. The actin fields are normalized via 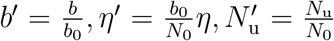. This way, the governing equations do not depend on the quantities *N*_0_ and *b*_0_ anymore:

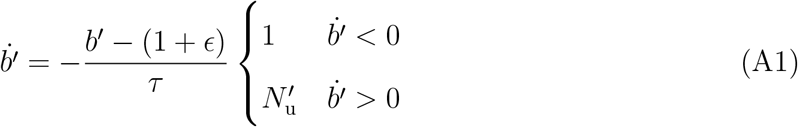

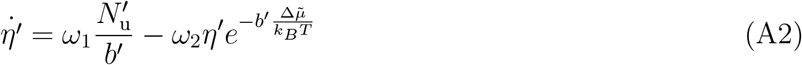

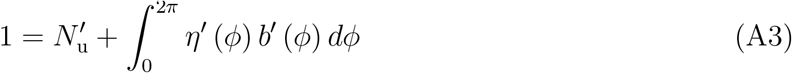

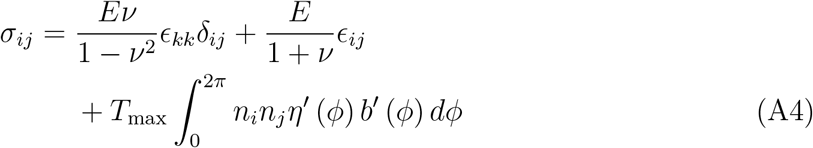

In the last expression *ϵ*_*ij*_ and *σ*_*ij*_ already denote the two-dimensional strain and stress tensors.

## Appendix B: Sensitivity of the Actin Ring to acto-myosin contractility

**FIG. 7:**
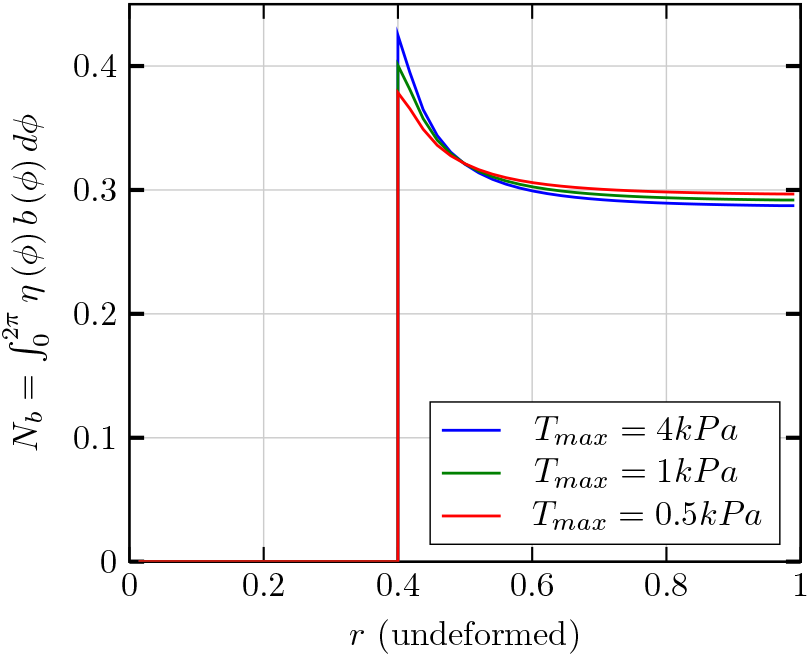
To check the influence of the contractile tension *T*_max_ the radial profiles of bound actin achieved in the steady state (compare figure 5b) are plotted for different values of *T*_max_. It can be seen that even if the contractile tension is decreased by a factor of 8 a significant actin ring develops. This is consistent with the experimental observation that the actin ring also develops after the addition of blebbistatin which is known to decrease acto-myosin contractility, as discussed in the section III A.

## ACKNOWLEDGMENTS

This work was funded by the Deutsche Forschungsgemeinschaft (DFG, German Research Foundation) under project number 546825574. B.B. acknowledges support from the Biological Physics study program of the Elite Network of Bavaria. This publication was funded by the University of Bayreuth Open Access Publishing Fund. Calculations were performed using the festus-cluster of the Bayreuth Centre for High Performance Computing funded by the Deutsche Forschungsgemeinschaft (DFG, German Research Foundation) under project number 523317330.

## DATA AVAILABILITY

The simulation code and resulting data used to obtain the results of this work are available at Zenodo under DOI 10.5281/zenodo.22806150.

## Footnotes

1 This technique measures the local cytoplasmic viscosity inside the cell using rotating wires with lengths *L* ≈ 4*µm* and comparably small rotation speeds 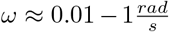. As this length scale is relatively close to the size of the nucleus we expect this viscosity to be a good approximation for the viscosity experienced by the nucleus.

